# Integrated single-cell and bulk transcriptomic analyses of lysine isonicotinylation-related gene expression in diabetic foot ulcer healing

**DOI:** 10.64898/2026.11.26.769490

**Authors:** Zijun Ye, Yinghao Wu, Chenhao Ye, Bohuai Ning, Lin Yang, Yiheng Chen

**Affiliations:** Department of Hand and Microsurgery, The Second Affiliated Hospital and Yuying Children’s Hospital of Wenzhou Medical University, Wenzhou, Zhejiang, China; Department of Pediatric Respiratory Medicine, National Key Clinical Specialty of Pediatric Respiratory Medicine, Institute of Pediatrics, The Second Affiliated Hospital, Yuying Children’s Hospital of Wenzhou Medical University, Wenzhou, Zhejiang, China; Department of Orthopedics, People’s Hospital of Jingning She Autonomous County, Lishui, Zhejiang, China

## Abstract

Diabetic foot ulcers have heterogeneous healing outcomes. We integrated public single-cell and bulk RNA-sequencing datasets to identify cell-type-specific lysine isonicotinylation (Kinic)-related transcriptional patterns and exploratory candidates associated with healing. The atlas comprised 56,568 cells across 14 cell types, with Kinic-related signature scores differing by cell type and healing group. Intersecting bulk differentially expressed genes, single-cell high-versus-low-score differentially expressed genes, and coexpression module genes prioritized LDHB, ISG15, and HAUS1 as exploratory candidates. Seven subtypes were identified among healing-enriched fibroblasts. ISG15-associated enrichment highlighted interferon and inflammatory responses. Simulated deletion of each candidate reduced mean similarity to a non-Healer reference state. Docking identified LDHB–ZINC250K_0191530 as the lowest-scoring evaluated pair, while a single 100-ns simulation showed displacement of the ligand from its initial pose. These analyses prioritize LDHB, ISG15, and HAUS1 as exploratory candidates and characterize their cellular context in diabetic foot ulcer healing. Their expression patterns within healing-enriched fibroblasts provide clues to the cellular and molecular differences associated with healing outcomes.

**Author summary:** Why do some foot ulcers in people with diabetes heal while others remain open? We examined gene activity in cells from ulcers with known healing outcomes. We focused on genes previously linked to a chemical change that can occur on proteins. Gene activity gives us an indirect way to study this question. Measuring the chemical change itself would require a different experiment. We found that a group of cells that help build and repair connective tissue showed higher combined activity of these genes in ulcers that healed. Two types of immune cells showed the opposite pattern. Comparing whole tissue with individual cells also highlighted three genes for closer study. One of them, ISG15, was associated with inflammatory signals. The whole-tissue comparison involved only thirteen samples, so the three genes are starting points for investigation. Our findings point to a focused question for future work: whether the differences we saw in repair cells recur across individual patients and accompany changes in the proteins themselves.

## Introduction

Diabetic foot ulcers (DFUs) are a disabling complication of diabetes that can require prolonged care and lead to amputation. Healing outcomes vary, suggesting differences in the cellular processes that sustain repair. Single-cell studies have identified healing-associated fibroblasts with inflammatory and extracellular matrix-remodeling features, together with distinct macrophage states [1,2]. Understanding how these wound-cell populations differ between healing and nonhealing ulcers may help identify processes associated with successful repair.

Lysine isonicotinylation (Kinic) was identified as a histone acylation mark induced by isoniazid and its metabolites. Jiang et al. linked this modification to increased chromatin accessibility and transcription [3]. Zhou et al. subsequently described an 84-gene Kinic-associated set assembled from GeneCards and published literature for transcriptomic analysis in hepatocellular carcinoma [4]. This work provides a rationale for examining related gene-expression patterns in other disease settings, although the biochemical relevance of such patterns must be assessed in the tissue under study.

Previous DFU studies have characterized healing-associated fibroblasts and implicated their inflammatory and matrix-remodeling programs in the wound microenvironment [1,2]. Whether Kinic-related transcriptional patterns differ between healing and nonhealing ulcers, and which cell populations account for those differences, remains unclear. Examining these patterns within defined populations could connect tissue-level expression differences with the cellular states involved in repair. Bulk RNA sequencing captures aggregate tissue expression, while single-cell analysis helps localize those signals and assess cellular heterogeneity.

We combined bulk and single-cell transcriptomes to compare gene expression between healing and nonhealing DFUs. We aimed to characterize cell-type-specific Kinic-related transcriptional patterns and prioritize candidate genes associated with healing status. These analyses may provide clues to the cellular and molecular differences accompanying heterogeneous healing outcomes.

## Materials and Methods

### Data acquisition and collection of Kinic-related genes

We obtained expression data and the associated sample annotations through the Gene Expression Omnibus (GEO). The single-cell datasets comprised eight DFU debridement samples from GSE231643 (five Healer and three non-Healer) [2] and 14 foot-skin samples from patients with DFUs in GSE165816 (nine Healer and five non-Healer) [1]. The bulk RNA-sequencing dataset comprised 13 ulcer-tissue samples from GSE134431 (seven Healer and six non-Healer) [5]. Healing-status labels were retained from the source studies. Gene-set scoring used the 84-gene Kinic-related input list supplied with the analysis files and provided in S1 Data. Zhou et al. describe the assembly of an 84-gene Kinic-associated set from GeneCards and published literature [4]. Table 1 summarizes the datasets, and S1 Data lists the included sample identifiers.

**Table 1.** Characteristics of the transcriptomic datasets included in this study. Counts represent samples included in the present analysis. Healer and non-Healer labels follow the original study annotations. DFU, diabetic foot ulcer; GEO, Gene Expression Omnibus; RNA-seq, RNA sequencing.

| **Dataset** | **Data type** | **Tissue source** | **Healer, n** | **non-Healer, n** | **Total, n** | **GEO platform** |
|---|---|---|---|---|---|---|
| GSE231643 | Single-cell RNA-seq | DFU debridement tissue | 5 | 3 | 8 | GPL18573 |
| GSE165816 | Single-cell RNA-seq | Foot skin from patients with DFUs | 9 | 5 | 14 | GPL24676 |
| GSE134431 | Bulk RNA-seq | DFU tissue | 7 | 6 | 13 | GPL18573 |

### Quality control and integration of single-cell RNA-sequencing data

We applied dataset-specific quality-control thresholds in Seurat (version 5.3.0) [6]. GSE231643 cells were retained if they had 500–30,000 unique molecular identifiers (UMIs), 200–6,000 detected genes, and ≤10% mitochondrial UMIs. For GSE165816, these thresholds were 500–15,000 UMIs, 200–4,000 detected genes, and ≤15% mitochondrial UMIs. DoubletFinder [7] was run separately for each sample using the first 20 principal components and pN = 0.25. For each sample, pK was selected at the maximum BCmetric, and the expected doublet number (nExp) was set to 5% of the input cell count. Predicted doublets were removed before downstream analysis. Cell counts at each filtering step are summarized.

Following doublet removal and re-normalization, the 2,000 highly variable genes selected by FindVariableFeatures were used for principal component analysis. Inspection of the elbow plot supported retaining the first 15 components. Harmony (version 1.2.3) was applied to reduce batch-related variation in the low-dimensional representation [8]. Sample mixing was assessed qualitatively using Uniform Manifold Approximation and Projection (UMAP) plots colored by sample before and after integration.

### Cell clustering and cell-type annotation

Integrated cells were clustered in Seurat at a resolution of 1.0. Cluster markers were identified with FindAllMarkers using the Wilcoxon rank-sum test, with min.pct set to 0.25 and logfc.threshold set to 0.25. Manual annotation drew on the top 10 markers in each cluster, marker information from Choi et al. [2], and CellMarker 2.0 [9]. UMAP plots displayed the annotated populations. Cell-type proportions were calculated from pooled cell counts within each healing group and displayed in an alluvial plot. A dot plot of representative markers supported the cell-type assignments.

### Kinic-related gene signature scoring

UCell [10] was used to score the Kinic-related signature in individual cells. Genes from the supplied Kinic-related list that were present in the expression matrix were included in scoring. Within each annotated cell type, scores were compared between Healer and non-Healer cells using Wilcoxon rank-sum tests. Unadjusted P < 0.05 defined exploratory cell-level differences. Cells were then assigned to High and Low groups using the median score across all annotated cells. These assignments were mapped onto the UMAP embedding, and the proportions of High- and Low-score cells were calculated for each cell type.

### High-dimensional weighted gene co-expression network analysis

Co-expression networks were constructed with high-dimensional weighted gene co-expression network analysis (hdWGCNA) [11] using genes detected in at least 5% of cells. Cells were grouped by annotated cell type and Kinic score group (High or Low), pooling cells from different samples within each group. Metacells were formed from 25 nearest neighbors. Network topology guided selection of the soft-thresholding power. Modules were defined using topological overlap and dynamic tree cutting, with a minimum size of 100 genes, a merging threshold of 0.2, and deepSplit = 4. Pearson correlation coefficients were used to assess the relationships between module eigengenes and Kinic score groups (High versus Low), with P < 0.05 as the criterion for statistical significance. Genes from the selected module were used in the subsequent intersection analysis.

### Differential expression analysis and candidate gene identification

Bulk RNA-sequencing counts from GSE134431 were compared between Healer and non-Healer samples using DESeq2 (version 1.48.2) [12]. Bulk differentially expressed genes (DEGs-1) were defined using |log₂FC| > 0.5 and unadjusted P < 0.05. These nominal thresholds were adopted after exploratory relaxation of the initial screening criteria to obtain an overlap with the other gene sets. For the single-cell comparison, FindMarkers in Seurat applied the Wilcoxon rank-sum test to Kinic High versus Low cells, pooling all annotated populations. Single-cell differentially expressed genes (DEGs-2) met unadjusted P < 0.05 and |log₂FC| > 0.25. The intersection of DEGs-1, DEGs-2, and the selected coexpression module defined the exploratory candidates. Overlaps were plotted with ggvenn (version 0.1.10) [13].

### Gene–gene interaction network analysis

GeneMANIA [14] was queried with the candidate genes obtained from the preceding screen to construct a gene–gene interaction (GGI) network. The network was expanded to include 20 additional functionally related genes. Edges represented database evidence for physical interactions, co-expression, predicted associations, and other relationships.

### Gene set enrichment analysis

Gene set enrichment analysis (GSEA) [15] was conducted for the candidate genes in the 13 bulk RNA-sequencing samples from GSE134431. For each candidate, genes were ranked by decreasing Spearman correlation with its expression across samples. Analysis used clusterProfiler (version 4.16.0) [16] and the human MSigDB Hallmark collection (h.all.v2025.1.Hs.symbols.gmt) [17]. Gene sets were retained when Benjamini–Hochberg-adjusted P was < 0.05 and the normalized enrichment score (NES) met |NES| > 1. The five retained sets with the highest absolute NES were plotted for each candidate.

### Immune infiltration analysis

MCP-counter (version 1.2.0) [18] estimated abundance scores for eight immune and two stromal populations in the 13 selected GSE134431 samples (7 Healer and 6 non-Healer). For heatmaps, scores were Z-standardized across samples within each population. Between-group differences were tested on the original scores using Mann–Whitney U tests. Benjamini– Hochberg adjustment covered the ten population comparisons, and both nominal and adjusted P values were reported. Populations with nominal between-group P < 0.05 were selected for exploratory correlation analysis. Spearman correlations between their abundance scores and candidate-gene expression were calculated across all 13 samples. Correlation P values were reported without adjustment.

### Single-cell expression analysis of candidate genes

Uniform Manifold Approximation and Projection (UMAP) feature plots displayed candidate-gene expression in the integrated single-cell dataset. Within cell types with differing Kinic-related signature scores between healing groups, candidate expression was compared using cell-level Wilcoxon rank-sum tests. Adjusted P < 0.05 defined exploratory expression differences. Z-score heatmaps summarized average expression by cell type and healing group. Cell types showing differences for all three candidates were considered for follow-up, with pooled cell proportions across healing groups and population size used to select the population for subclustering.

### Cell–cell communication analysis

CellChat (version 2.1.2) [19,20] and the human CellChatDB were used to infer cell–cell communication from the data layer of the RNA assay. Networks were constructed separately for Healer and non-Healer cells, with annotated cell types defining the communication groups. Interactions were excluded if either cell group contained fewer than 10 cells (min.cells = 10), and the P-value cutoff was 0.05. We compared total interaction counts and aggregated strengths between the networks. Circular plots showed interactions among cell types, while relative information flow described communication at the pathway level. Outgoing and incoming signaling strengths across all pathways were examined to identify potential sender and receiver roles.

### Subclustering and pseudotime analysis of the selected fibroblast population

The fibroblast population selected through the preceding expression and composition analysis was reanalyzed in Seurat [6] using 2,000 highly variable genes, the first 10 principal components, and a clustering resolution of 0.4. Cluster markers were identified with FindAllMarkers using min.pct = 0.5, logfc.threshold = 0.25, and test.use = “auc”. Marker profiles and CellMarker 2.0 [9] guided subtype annotation. UMAP plots displayed the subtypes, and dot plots summarized representative marker expression. Subtype proportions were calculated from pooled fibroblast counts within each healing group.

Trajectory analysis used 3,000 selected fibroblasts, comprising 1,500 cells randomly sampled from each healing group. Monocle 2 (version 2.26.0) [21] inferred trajectories under a negative binomial expression model. Genes detected in at least 10 cells (min_expr = 0.1) were retained; ordering genes additionally required mean_expression ≥ 0.1 and dispersion_empirical ≥ dispersion_fit. DDRTree reduced the data to two components.

Following the initial ordering, the state with the highest CytoTRACE scores [22] was chosen as the root. A second run of orderCells assigned the final pseudotime. Spearman correlation described the relationship between CytoTRACE scores and pseudotime. Candidate-gene expression was plotted along the inferred trajectory.

### In silico gene perturbation analysis

Candidate-gene knockout was simulated in the selected fibroblast population with scTenifoldKnk (version 1.0.3) [23]. Gene regulatory networks containing 5,000 genes were built from 10 resampling runs of up to 500 cells. For each knockout, the target gene’s outgoing connections were removed before manifold alignment of the original and perturbed networks. Excluding the perturbed gene itself, genes meeting both Benjamini–Hochberg-adjusted P < 0.05 and |Z| ≥ 2 were selected for enrichment analysis. Gene Ontology (GO) and Kyoto Encyclopedia of Genes and Genomes (KEGG) enrichment used Entrez Gene identifiers and clusterProfiler [16]. Terms with adjusted P < 0.05 were retained, and up to ten per category were displayed in increasing order of adjusted P.

Geneformer-V2-316M [24,25] was applied to selected fibroblasts without further fine-tuning, with deletion-type perturbations evaluated in Healer selected fibroblasts. Raw counts were tokenized after gene symbols were mapped to Ensembl IDs, using a 4,096-token limit. Healer and non-Healer cells defined the start and goal states, respectively. Each group was required to contain at least 10 cells of the selected type. Candidate genes had to map to Ensembl IDs and occur in the input token sequences of Healer cells. In the embedding analysis, each state was represented by the mean last-layer classification-token (CLS) embedding. Each candidate gene was deleted individually from the token encoding. Final runs included up to 1,000 cells and used a forward batch size of 25. Shift_to_goal_end measured the change in cosine similarity to the non-Healer reference state, with negative values denoting reduced similarity. Mean shifts were used to summarize the predicted responses.

### GraphBAN-based virtual screening of candidate compounds

GraphBAN [26] screened 249,455 compounds from the ZINC250K library against the proteins encoded by the candidate genes. Inputs comprised compound structures encoded as simplified molecular-input line-entry system (SMILES) strings and target protein sequences, together with weights pretrained on BindingDB. Predictions were generated in 10 computational batches and ranked separately for each target. Score distributions and counts above illustrative thresholds of 0.50, 0.60, and 0.75 were summarized to describe the predictions. Selection for physicochemical filtering used the 500 highest-scoring compounds for each target. The compound inputs and target-specific predictions are provided in S2–S4 Data.

For each target, the 500 highest-scoring compounds were filtered by molecular weight <500 Da, 1 ≤ logP ≤ 5, fewer than five hydrogen-bond acceptors, fewer than five hydrogen-bond donors, and topological polar surface area <140 Å². Compounds that passed these criteria were combined and deduplicated by compound identifier.

### Molecular docking

Selected compounds remaining after physicochemical filtering were docked to the proteins encoded by the candidate genes using CB-DOCK2 [27]. Ligand structures were obtained from PubChem in SDF format, and receptor structures were retrieved from RCSB PDB and AlphaFold DB in PDB format. Table 2 summarizes the receptor models used for docking.

**Table 2.** Receptor models used for molecular docking.

| **Protein** | **Receptor representation** |
|---|---|
| LDHB | Two protein chains retained in the docking model |
| ISG15 | Model containing the C78S substitution |
| HAUS1 | Fragment spanning residues 77–278 |

Docking scores were recorded in kcal/mol. PyMOL (version 3.1) [28] visualized predicted poses and local ligand–residue contacts, retaining residue numbering from the receptor files.

### Molecular dynamics simulation

The compound–protein complex with the lowest docking score was selected for a 100-ns molecular dynamics simulation in GROMACS (version 2025.2) [29]. The protein was modeled with the AMBER99SB-ILDN force field [30]. The ligand was assigned AM1-BCC partial charges [31] with a specified net charge of +1. Sobtop (version 2026.1.16) [32] generated ligand topology and parameters from GAFF atom types and preset parameters [33]. The complex was solvated in a cubic TIP3P water box [34] with a 10 Å boundary distance and neutralized with sodium and chloride ions. The system then underwent energy minimization, followed by NVT equilibration (100 ps, 300 K) and NPT equilibration (100 ps, 1 bar).

Trajectories were analyzed with GROMACS [29]. Backbone root-mean-square deviation (RMSD) was calculated after backbone alignment. Additional measures comprised atom-wise root-mean-square fluctuation (RMSF), protein–ligand hydrogen bonds, radius of gyration (Rg), and solvent-accessible surface area (SASA). A free-energy landscape was constructed from backbone RMSD and Rg with 80 × 80 bins and kT = 2.5 kJ/mol; empty bins were assigned the minimum nonzero density. In a separate post-processing analysis, ligand RMSD was reassessed using GROMACS (version 2025.1) after whole and nojump processing. Mass-weighted fitting used the N, Cα, and C backbone atoms from both protein chains. RMSD was calculated for all 35 ligand atoms relative to the production-run input structure, retaining all 2,501 frames without smoothing. Ligand poses at 0, 90.96, 91.80, 92.12, and 100 ns were compared in the protein-aligned reference frame without ligand self-alignment.

### Ethics statement

This study reanalyzed publicly available transcriptomic datasets. No participants were recruited and no new human specimens or animal experiments were involved in the present analysis. No new ethics approval was sought for this secondary analysis. The source studies and their datasets are cited in the manuscript.

### Use of generative artificial intelligence in manuscript preparation

We used ChatGPT and Codex (OpenAI) to assist with manuscript drafting, language editing, consistency checks, preparation of the plain-language Author summary, and formatting for submission.

## Results

### Construction and annotation of an integrated single-cell transcriptomic atlas of diabetic foot ulcers

GSE231643 and GSE165816 were integrated according to the study workflow (Fig 1A), initially yielding 69,352 cells. Dataset-specific quality control retained 59,546 cells. Removal of 2,978 predicted doublets left 56,568 cells for analysis (Fig 1B). UMAP inspection suggested greater mixing of cells from different samples after Harmony integration (Fig 1C).

**Fig 1.**
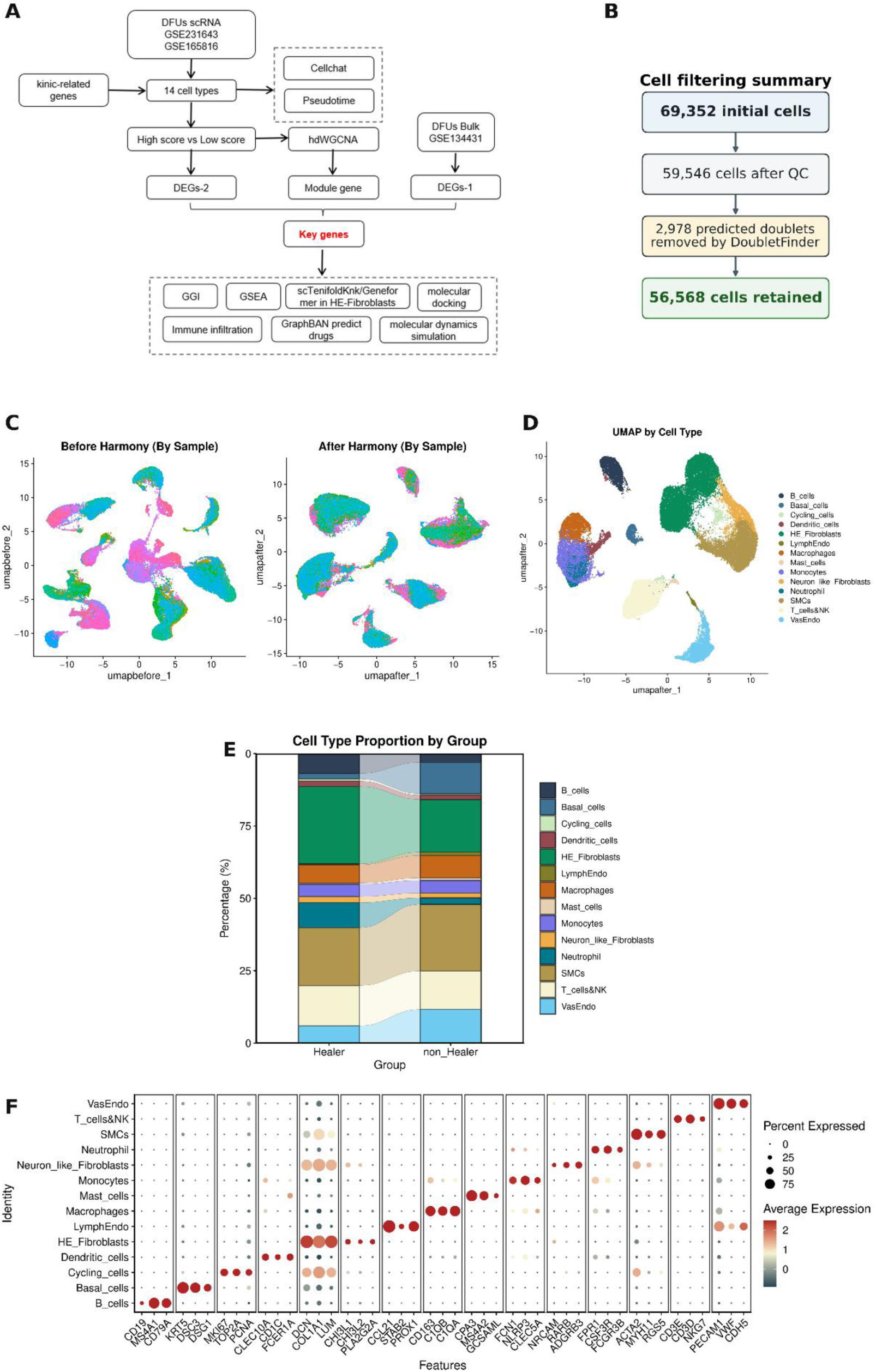
Construction and annotation of an integrated single-cell transcriptomic atlas of diabetic foot ulcers. (A) Study workflow. (B) Cell counts retained after quality-control filtering and doublet removal. (C) UMAP plots before and after Harmony integration, colored by sample. (D) UMAP of 56,568 cells colored by the 14 annotated populations. (E) Cell-type proportions based on pooled counts within the Healer and non-Healer groups. Bands connect the same cell categories across groups and do not represent cell transitions. (F) Representative marker expression across annotated populations. Dot size denotes the percentage of cells expressing each gene; color denotes scaled average expression.

Unsupervised clustering identified 26 clusters, which were assigned to 14 major cell populations (Fig 1D). The largest populations were healing-enriched fibroblasts (HE-Fibroblasts; 13,456 cells, 23.8%) and smooth muscle cells (11,893 cells, 21.0%). In pooled cell counts, HE-Fibroblasts accounted for a larger proportion of Healer cells, whereas basal and vascular endothelial cells were proportionally more abundant in non-Healer cells (Fig 1E). Representative marker profiles supported the annotations (Fig 1F).

### Cell type-specific patterns of Kinic-related gene signature scores

The Kinic-related signature was scored using the 78 of 84 genes present in the expression matrix. Exploratory cell-level comparisons detected differences between Healer and non-Healer cells in 10 of 14 populations (nominal P < 0.05). Healer HE-Fibroblasts had higher scores, whereas Healer monocytes and neutrophils had lower scores (Fig 2A). The median score across all cells separated 28,286 high-score cells from 28,282 low-score cells (Figure 2B). The proportion of high-score cells ranged from 12.8% in vascular endothelial cells to 80.6% in monocytes (Fig 2C).

**Fig 2.**
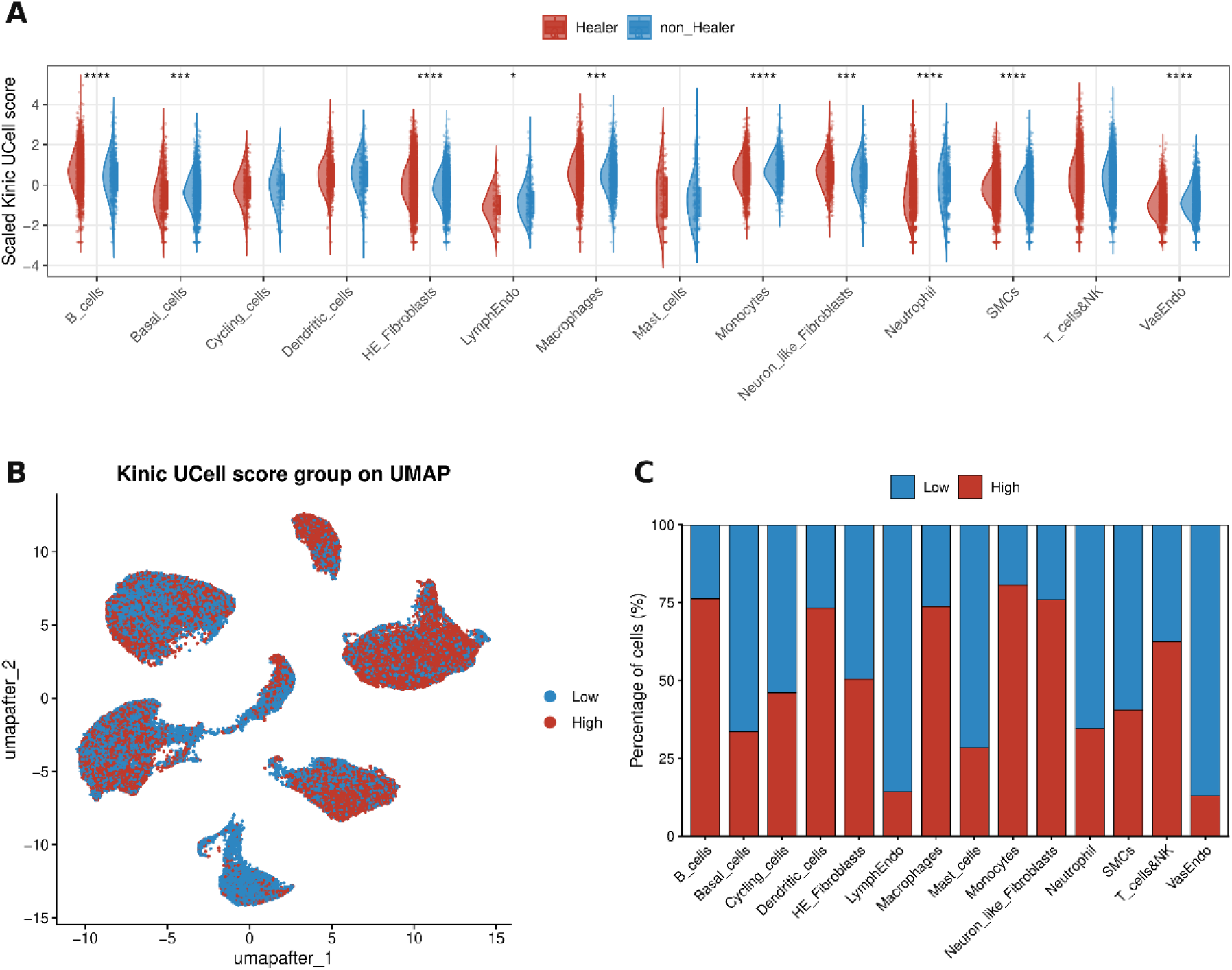
Cell type-specific patterns of Kinic-related gene signature scores. (A) Scaled UCell score distributions in Healer and non-Healer cells across the annotated populations. Significance annotations represent nominal P values from cell-level Wilcoxon rank-sum tests. One, two, three, and four asterisks indicate P < 0.05, 0.01, 0.001, and 0.0001, respectively. (B) UMAP visualization of high- and low-score cells defined using the median score across all cells. (C) Percentages of high- and low-score cells within each annotated population.

### Identification of a Kinic-associated co-expression module and candidate genes

At a soft-thresholding power of 12, the hdWGCNA network had a scale-free topology fit of R² = 0.825 (Fig 3A). Hierarchical clustering identified 12 co-expression modules, excluding the unassigned genes in the gray group (Fig 3B). Descriptive comparison of the module profiles showed higher MEs-1 values in Kinic low-score cells than in high-score cells (Fig 3C). We carried MEs-1 forward for exploratory screening and intersected its 768 genes with the bulk and single-cell differential expression results.

**Fig 3.**
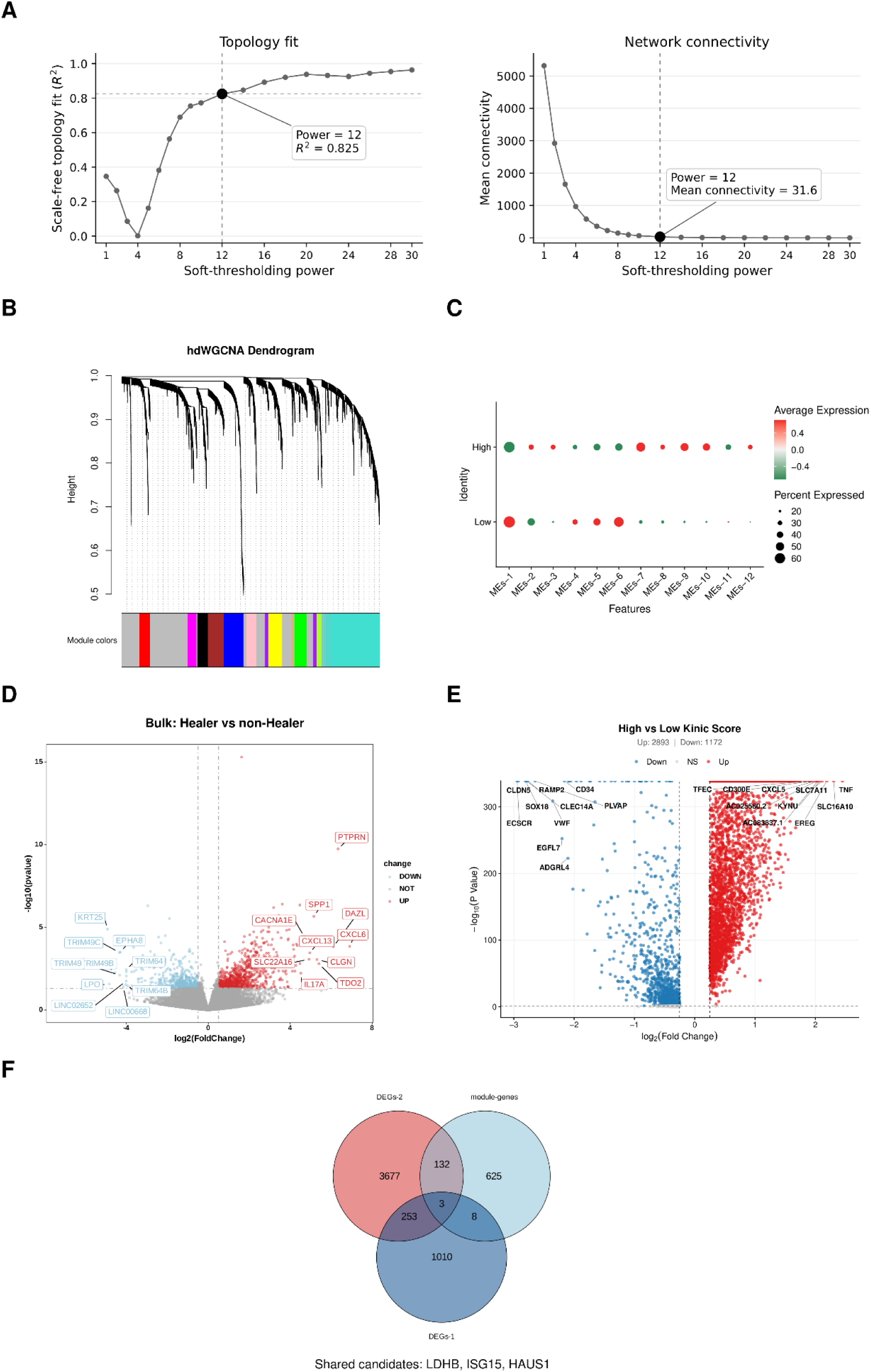
Identification of a Kinic-associated co-expression module and candidate genes. (A) Scale-free topology fit and mean connectivity across tested soft-thresholding powers, with the selected power of 12 highlighted. (B) Gene clustering dendrogram and module assignments; gray denotes unassigned genes. (C) Module profiles in Kinic high- and low-score cells. Color represents the scaled average expression displayed by the module-profile plot, and dot size represents the percentage expressed. MEs-1 was used for subsequent exploratory screening. (D) Bulk differential expression between Healer and non-Healer samples at nominal P < 0.05 and |log₂FC| > 0.5. (E) Differential expression between Kinic high- and low-score cells at nominal P < 0.05 and |log₂FC| > 0.25. In the volcano plots in D and E, red and blue denote higher and lower expression, respectively, in the first named group; gray denotes genes that did not meet the selection criteria. (F) Venn diagram showing the intersection of DEGs-1, DEGs-2, and the 768 MEs-1 genes, identifying LDHB, ISG15, and HAUS1 as shared candidates.

Using the thresholds described in Methods, the bulk comparison yielded 1,274 DEGs-1 (815 with higher and 459 with lower expression in Healer than in non-Healer samples; nominal P < 0.05 and |log₂FC| > 0.5; Fig 3D). The comparison of Kinic high- and low-score cells yielded 4,065 DEGs-2, with 2,893 upregulated and 1,172 downregulated in high-score cells (nominal P < 0.05 and |log₂FC| > 0.25; Fig 3E). Intersecting these lists with the 768 MEs-1 genes identified LDHB, ISG15, and HAUS1 (Fig 3F). Their bulk adjusted P values were 0.291, 0.663, and 0.614, respectively. All three were carried forward as exploratory candidates.

### Interaction networks and functional enrichment of candidate genes

The GeneMANIA network connected LDHB, ISG15, and HAUS1 to 20 additional functionally related genes, with edges supported by physical interactions, co-expression, predicted associations, and other evidence categories (Fig 4C).

**Fig 4.**
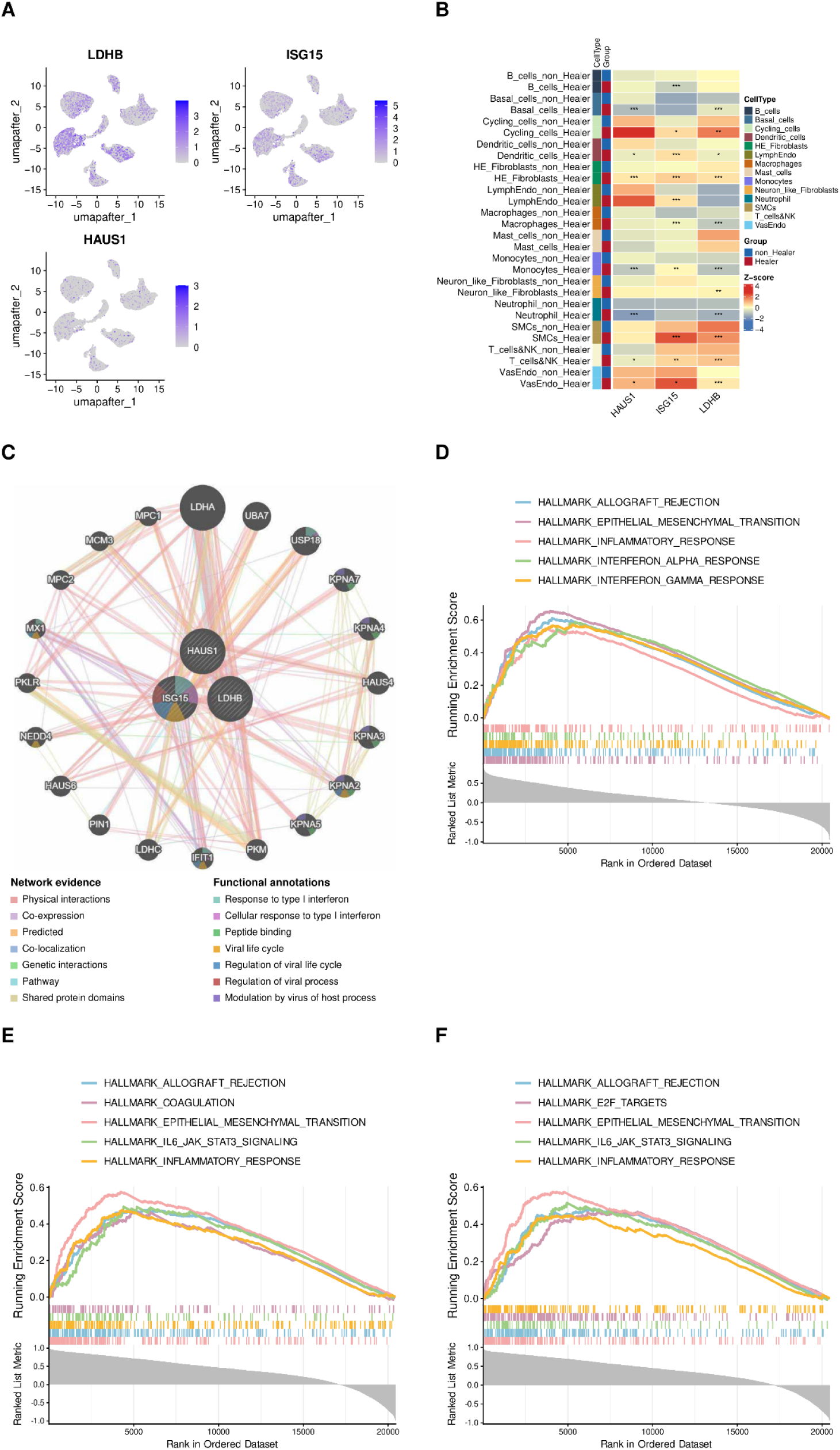
Expression patterns and functional characterization of the candidate genes. (A) UMAP feature plots of LDHB, ISG15, and HAUS1 expression, with a separate color scale for each gene. (B) Heatmap of Z-score-scaled average expression by cell type and healing group. Asterisks denote adjusted P values from cell-level Wilcoxon rank-sum tests comparing Healer and non-Healer cells within each type. One, two, and three asterisks indicate adjusted P < 0.05, 0.01, and 0.001, respectively. (C) GeneMANIA network of the three candidates and 20 additional functionally related genes. Edge colors denote evidence categories, and colored node segments denote functional annotations. (D–F) GSEA plots for ISG15 (D), HAUS1 (E), and LDHB (F). Each panel shows the five significant Hallmark sets with the highest absolute NES (adjusted P < 0.05 and |NES| > 1). Curves represent running enrichment scores; vertical ticks locate gene set members in the ranked list. The lower panel shows the ranking metric.

Hallmark GSEA detected 23, 20, and 21 significantly enriched gene sets for ISG15, HAUS1, and LDHB, respectively (adjusted P < 0.05 and |NES| > 1). Of the 23 ISG15-associated sets, 20 had positive and three had negative NES values. Epithelial–mesenchymal transition, allograft rejection, and inflammatory response ranked among the five leading sets in each analysis (Figures 4D–F). For ISG15, the other leading sets were interferon-α and interferon-γ responses (Fig 4D). For HAUS1, they were IL6–JAK–STAT3 signaling and coagulation (Fig 4E); for LDHB, they were IL6–JAK–STAT3 signaling and E2F targets (Fig 4F). All displayed sets had positive NES values.

### Immune-cell abundance and candidate-gene correlations in bulk tissue

MCP-counter profiled eight immune and two stromal populations in the 13 GSE134431 samples (Fig 5A). Neutrophil scores were higher in Healer samples (nominal P = 0.038; adjusted P = 0.383), but none of the ten population comparisons met adjusted P < 0.05 (Fig 5B). Across all samples, neutrophil scores correlated positively with HAUS1 (Spearman ρ = 0.86, P < 0.001), LDHB (ρ = 0.75, P = 0.003), and ISG15 expression (ρ = 0.66, P = 0.013) (Fig 5C). These exploratory correlations used unadjusted P values.

**Fig 5.**
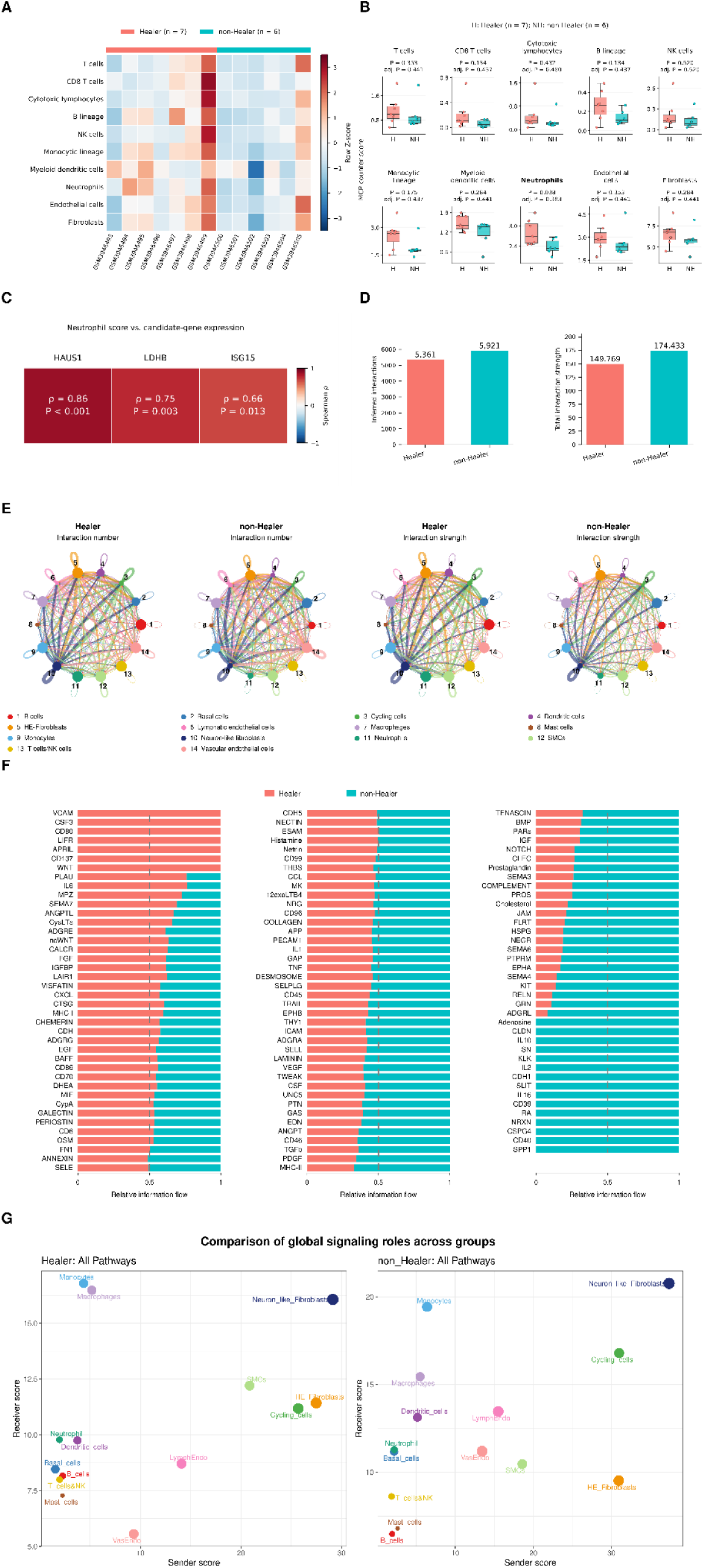
Immune microenvironment and intercellular communication patterns associated with DFU healing. (A) Heatmap of MCP-counter scores for eight immune and two stromal populations in 13 GSE134431 samples, Z-standardized across samples within each population. (B) Scores in Healer (n = 7) and non-Healer (n = 6) samples. Points denote individual samples, boxes show medians and interquartile ranges, and whiskers extend to the most extreme observations within 1.5 times the interquartile range. Nominal and Benjamini–Hochberg-adjusted P values are shown. (C) Spearman correlations between neutrophil scores and candidate-gene expression across all 13 samples, with correlation coefficients and unadjusted P values. (D) Total interaction counts and strengths inferred by CellChat for the Healer and non-Healer networks. (E) Circular networks of interaction counts and strengths among the 14 cell populations in each group. Numbered nodes refer to cell types in the shared key. (F) Relative information flow for 118 signaling pathways arranged in three columns. Each stacked bar represents the two groups’ relative contributions to the inferred information flow of a pathway. (G) Outgoing (sender) and incoming (receiver) signaling scores by cell population and healing group. CellChat comparisons are descriptive. H, Healer; NH, non-Healer.

### Single-cell expression of candidate genes and fibroblast selection

Expression of LDHB, ISG15, and HAUS1 varied across the integrated cell populations (Fig 4A). In HE-Fibroblasts and vascular endothelial cells, all three genes had higher average expression in Healer than in non-Healer cells (adjusted P < 0.05 in exploratory cell-level comparisons; Fig 4B). In the pooled cell composition, HE-Fibroblasts accounted for a larger proportion of Healer cells, whereas vascular endothelial cells accounted for a smaller proportion (Fig 1E). Together with the larger size of the HE-Fibroblast population, these observations motivated its selection for further analysis.

### Inferred cell–cell communication in the single-cell atlas

CellChat inferred 5,361 interactions in Healer and 5,921 in non-Healer cells, with total interaction strengths of 149.769 and 174.433, respectively (Fig 5D). The circular network plots showed differences across the 14 populations, with vascular endothelial connections visually less prominent in the Healer network (Fig 5E). Seven pathways, including VCAM and CSF3, had nonzero inferred information flow only in Healer cells. Conversely, 15 pathways, including SPP1 and CD40, had nonzero flow only in non-Healer cells (Fig 5F). Separately, the sender–receiver role analysis showed that HE-Fibroblasts had lower outgoing and higher incoming signaling scores in Healer than in non-Healer cells (Fig 5G). Network visualizations and sender–receiver scores summarize different aspects of inferred communication and were interpreted descriptively.

### Heterogeneity and pseudotemporal dynamics of HE-Fibroblasts

Re-clustering the 13,456 HE-Fibroblasts resolved seven transcriptional subpopulations: adventitial (Adventitial_Fib), extracellular matrix-associated (ECM_Fib), homeostatic (Homeostatic_Fib), inflammatory (Inflammatory_Fib), matrix (Matrix_Fib), myofibroblast-like (Myofibroblast_like_Fib), and repair-activated (Repair_activated_Fib) fibroblasts (Fig 6A). Representative markers supported these annotations (Fig 6B). In pooled counts, Inflammatory_Fib and Matrix_Fib cells constituted larger proportions of HE-Fibroblasts in Healer than in non-Healer cells (18.49% versus 9.50% and 12.32% versus 6.96%, respectively). Adventitial_Fib, Homeostatic_Fib, and Repair_activated_Fib cells showed the reverse pattern (Fig 6C).

**Fig 6.**
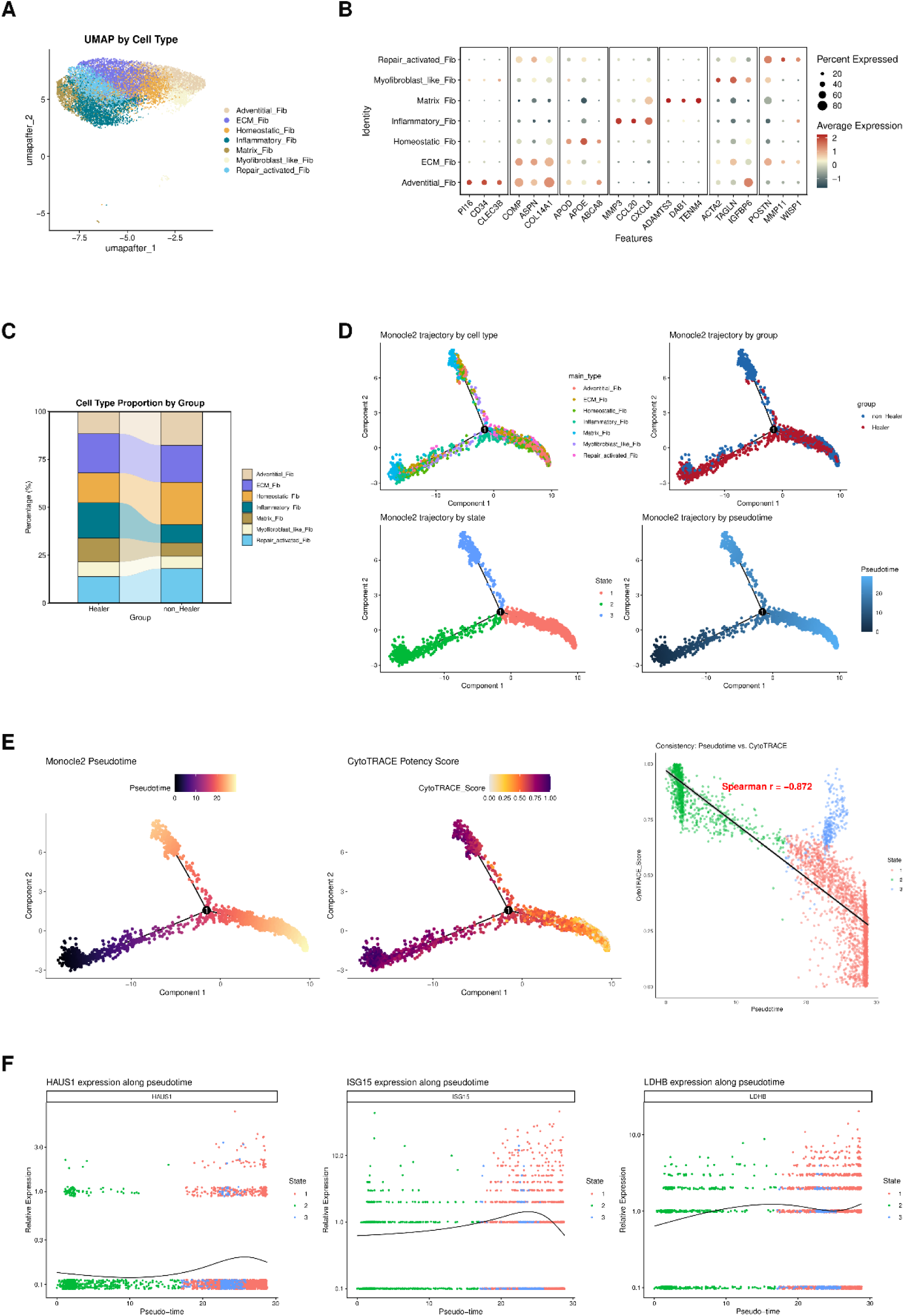
Heterogeneity and pseudotemporal dynamics of HE fibroblasts. (A) UMAP of seven transcriptional subpopulations within HE-Fibroblasts, labeled Adventitial_Fib, ECM_Fib, Homeostatic_Fib, Inflammatory_Fib, Matrix_Fib, Myofibroblast_like_Fib, and Repair_activated_Fib. (B) Representative marker expression across the seven subpopulations. Dot size denotes the percentage of expressing cells, and color denotes average expression. (C) Subpopulation proportions based on pooled HE-Fibroblast counts within each healing group. Each group was normalized to 100%, and differences were interpreted descriptively. (D) Monocle 2 trajectory of 3,000 HE-Fibroblasts, including 1,500 cells randomly sampled from each healing group. Cells are colored by subpopulation, healing group, model-derived trajectory state, and pseudotime. CytoTRACE scores guided root selection, and the numbered black circle marks the inferred branch point. (E) Monocle pseudotime and CytoTRACE scores along the trajectory, with their overall Spearman correlation. Because CytoTRACE guided root selection, this association is descriptive. (F) HAUS1, ISG15, and LDHB expression along inferred pseudotime. Cells are colored by trajectory state, and black curves show fitted trends.

A random sample of 1,500 HE-Fibroblasts from each healing group yielded a 3,000-cell trajectory with three states. The branch assigned to State 3 by the trajectory model predominantly contained non-Healer cells (Fig 6D). CytoTRACE scores remained relatively high in this branch despite an overall inverse correlation with pseudotime (Spearman ρ = −0.872; Fig 6E). State 3 denotes a model-derived branch state, and its assignment alone provides no evidence of terminal differentiation. The fitted expression of HAUS1 remained low, with a modest increase late in the trajectory. ISG15 expression increased and then fell near the endpoint; LDHB increased initially, declined, and rose again (Fig 6F).

### scTenifoldKnk and Geneformer-based virtual perturbation analysis

After simulated knockout of ISG15, HAUS1, and LDHB in HE-Fibroblasts, scTenifoldKnk predicted 63, 57, and 83 differentially regulated genes, respectively, excluding the perturbed gene itself (adjusted P < 0.05). Requiring |Z| ≥ 2 further narrowed these sets to 51, 34, and 75 genes (Fig 7A). Exploratory enrichment of the predicted responses to ISG15 deletion included extracellular matrix organization and extracellular matrix structural constituent terms. No KEGG pathway met adjusted P < 0.05 for LDHB (Fig 7B). These post-perturbation enrichment results are preliminary.

**Fig 7.**
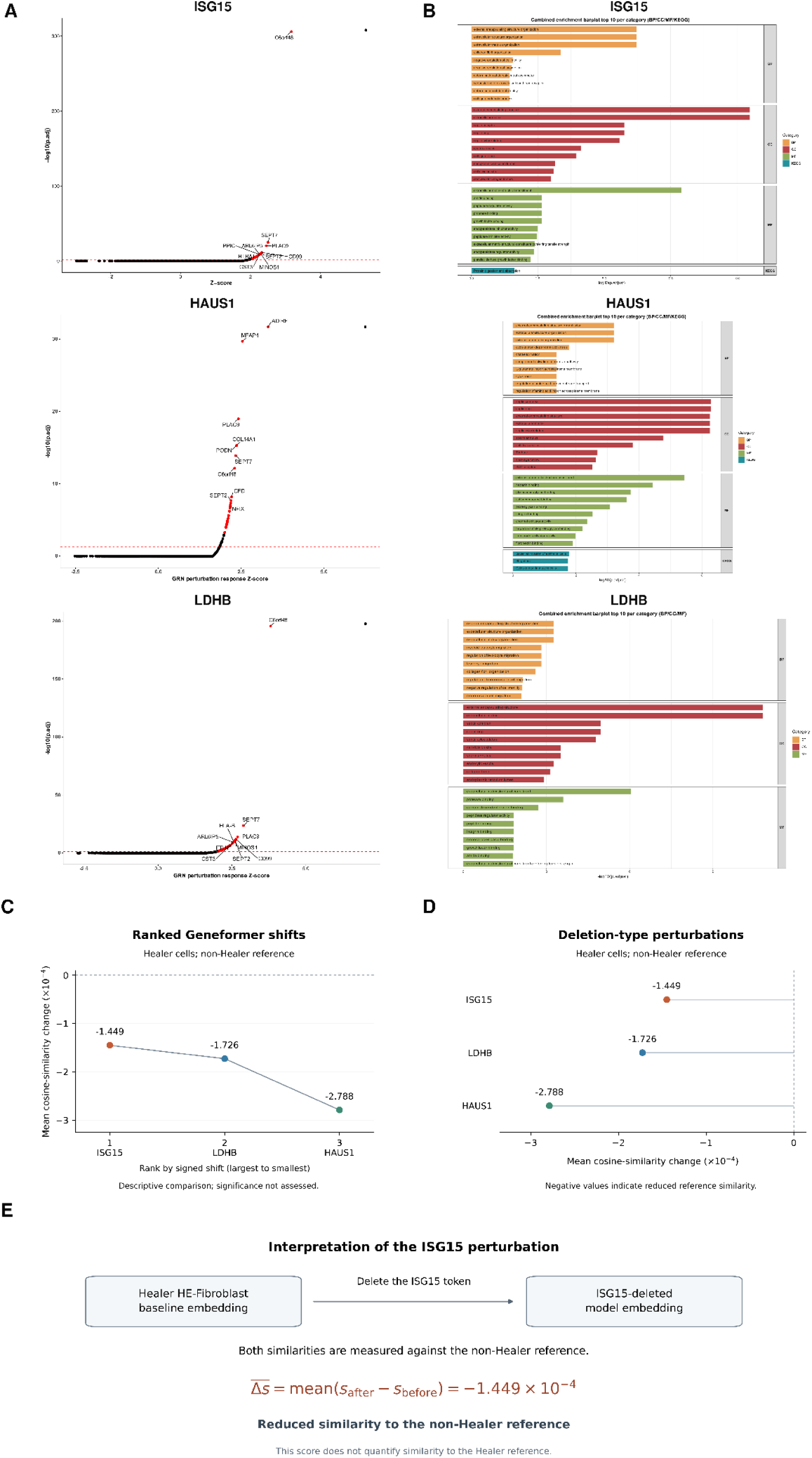
scTenifoldKnk and Geneformer-based virtual perturbation analysis of candidate genes in HE fibroblasts. (A) Predicted differential regulation after simulated knockout of ISG15, HAUS1, and LDHB. Horizontal axes show perturbation Z-scores; vertical axes show −log10(adjusted P). Red points denote genes meeting adjusted P < 0.05 and |Z| ≥ 2, excluding the perturbed gene itself. (B) Exploratory GO and KEGG enrichment of the predicted perturbed genes meeting adjusted P < 0.05 and |Z| ≥ 2, excluding the target gene. Up to ten significant terms per category are shown, with bar lengths representing −log10(adjusted P). BP, biological process; CC, cellular component; MF, molecular function. No KEGG pathway met adjusted P < 0.05 for LDHB. (C) The three tested genes ranked by mean cosine-similarity change (Shift_to_goal_end), from largest to smallest signed value. (D) Mean changes after deletion-type perturbations in Healer HE-Fibroblasts relative to the non-Healer reference state. Negative values indicate reduced reference similarity. (E) Schematic of the ISG15 perturbation. The arrow denotes deletion of the ISG15 token; baseline and perturbed embeddings are both compared with the non-Healer reference. Panels C–E show descriptive predictions without statistical significance testing.

Geneformer evaluated deletion-type perturbations in Healer HE-Fibroblasts relative to the non-Healer reference state. The mean changes in cosine similarity (Shift_to_goal_end) were −1.45 × 10⁻⁴ for ISG15, −1.73 × 10⁻⁴ for LDHB, and −2.79 × 10⁻⁴ for HAUS1 (Figures 7C, D). HAUS1 had the largest absolute mean change, and ISG15 had the smallest. All changes were negative, indicating reduced similarity to the non-Healer reference. The ISG15 result is shown schematically (Fig 7E).

### GraphBAN-based virtual screening and molecular docking

GraphBAN evaluated 249,455 ZINC250K compounds against ISG15, HAUS1, and LDHB. ISG15 score distributions were similar across the ten computational batches (Fig 8A). For ISG15, 2,420, 388, and 52 compounds reached the illustrative score thresholds of ≥0.50, ≥0.60, and ≥0.75, respectively. ZINC250K_0023529 ranked highest with a predicted interaction score of 0.900 (Fig 8B). Structures of the 12 highest-scoring ISG15 candidates are shown (Fig 8C). For each of the three targets, the top 500 compounds by target-specific score entered physicochemical filtering. This filtering retained 74 ISG15-, 32 HAUS1-, and 18 LDHB-associated candidates. Merging overlapping candidates left 93 unique compounds.

**Fig 8.**
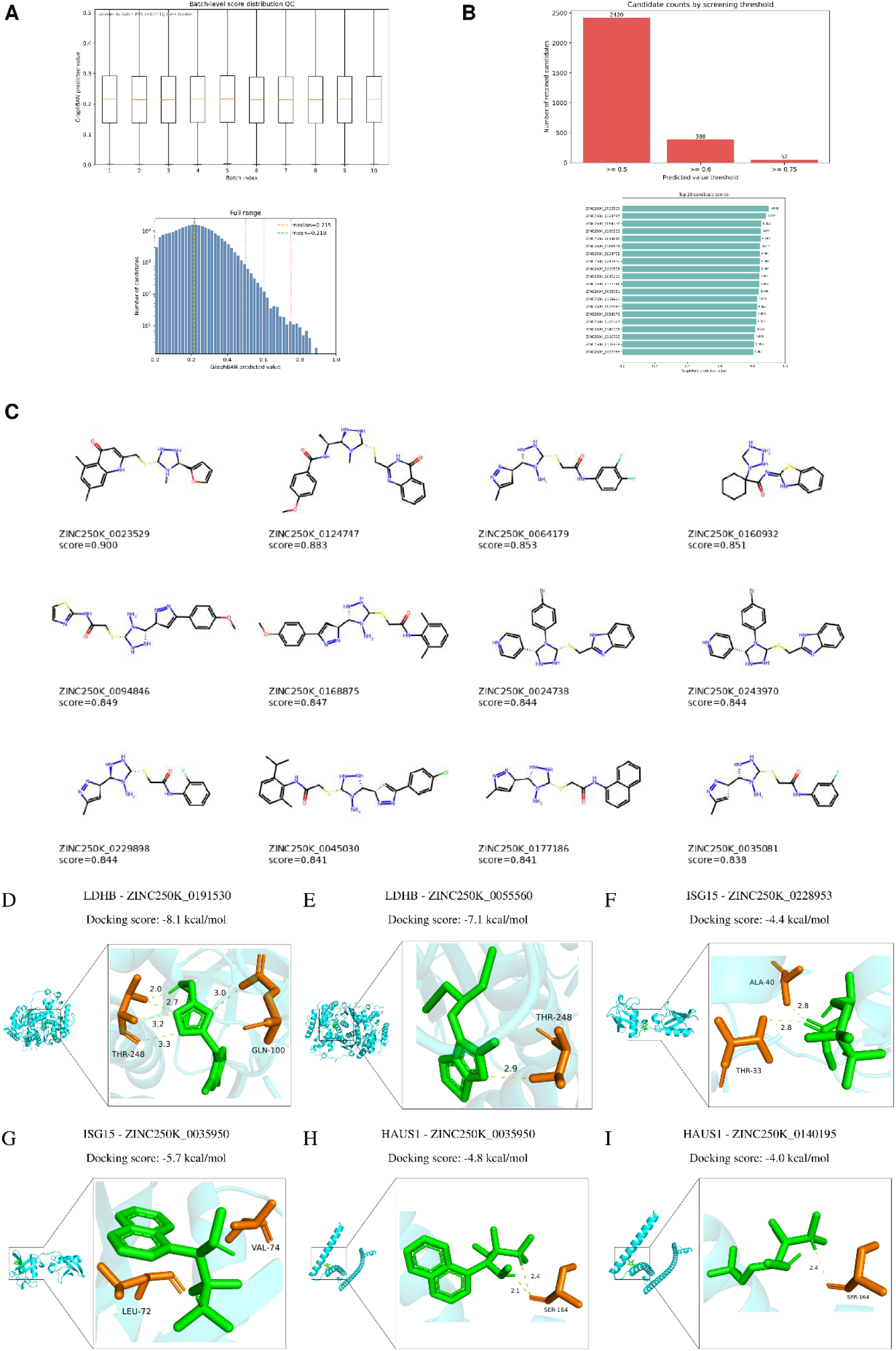
GraphBAN-based virtual screening and molecular docking of candidate compounds. (A) GraphBAN-predicted interaction scores for ISG15 across ten computational batches (upper) and the full candidate library (lower). The histogram count axis is logarithmic. (B) Compound counts at the indicated thresholds (upper) and the 20 highest-scoring ISG15 candidates (lower). (C) Two-dimensional structures of the 12 highest-scoring ISG15 candidates, labeled with compound identifiers and predicted scores. Rankings in B and C precede physicochemical filtering. (D, E) Predicted LDHB docking poses with ZINC250K_0191530 and ZINC250K_0055560, respectively. (F, G) Predicted poses of the ISG15 C78S receptor model with ZINC250K_0228953 and ZINC250K_0035950, respectively. (H, I) Predicted poses of the HAUS1 receptor fragment spanning residues 77–278 with ZINC250K_0035950 and ZINC250K_0140195, respectively. Receptors appear as cyan cartoons, ligands as green sticks, and selected receptor residues as orange sticks. Yellow dashed lines indicate the displayed interatomic distances (Å). Residue numbering follows the receptor files, and docking scores are given in kcal/mol.

Molecular docking evaluated six target–compound pairs comprising five unique compounds. LDHB docking scores were −8.1 kcal/mol for ZINC250K_0191530 and −7.1 kcal/mol for ZINC250K_0055560 (Figures 8D, E). ISG15 scores were −4.4 kcal/mol for ZINC250K_0228953 and −5.7 kcal/mol for ZINC250K_0035950 (Figures 8F, G). HAUS1 scores were −4.8 kcal/mol for ZINC250K_0035950 and −4.0 kcal/mol for ZINC250K_0140195 (Figures 8H, I). With the lowest docking score among the six pairs, LDHB–ZINC250K_0191530 was selected for a 100-ns molecular dynamics simulation.

### Molecular dynamics simulation of the LDHB–ZINC250K_0191530 complex

The LDHB–ZINC250K_0191530 complex was examined over a 100-ns molecular dynamics trajectory. After backbone alignment, backbone RMSD averaged 0.298 nm during 50–100 ns, with a range of 0.247–0.358 nm (Fig 9A). Atom-wise RMSF was below 0.40 nm for 95.6% of the analyzed atoms (Fig 9B). Protein–ligand hydrogen-bond counts ranged from zero to three; 227 of 501 sampled frames (45.3%) contained no hydrogen bonds (Fig 9C).

**Fig 9.**
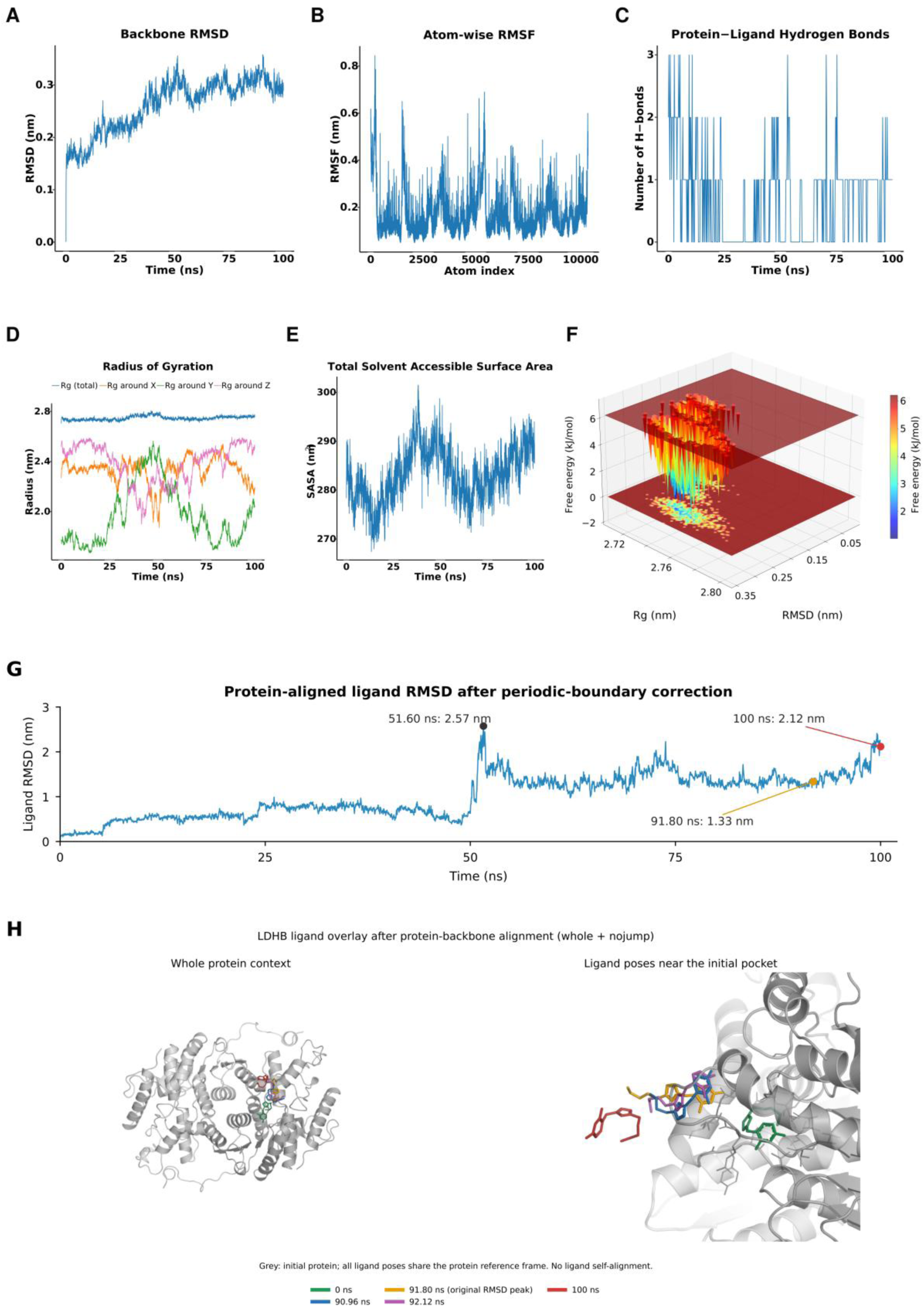
Molecular dynamics simulation of the LDHB–ZINC250K_0191530 complex. (A) LDHB backbone RMSD relative to the reference structure after least-squares alignment to the backbone. (B) Atom-wise RMSF profile, with atom indices shown on the horizontal axis. (C) Numbers of protein–ligand hydrogen bonds across 501 sampled frames over the 100-ns trajectory. Time is presented in nanoseconds. (D) Total radius of gyration (Rg) and radii of gyration about the x-, y-, and z-axes over the 100-ns trajectory. (E) Total solvent-accessible surface area (SASA) over time. (F) Free-energy landscape projected onto backbone RMSD and radius of gyration, constructed using 80 × 80 bins and kT = 2.5 kJ/mol. Empty bins were assigned the minimum nonzero density before conversion to relative free energy. Values in these bins therefore do not represent sampled conformations. The landscape summarizes conformational sampling within the analyzed trajectory and does not represent protein–ligand binding free energy. (G) Ligand RMSD after whole/nojump processing and protein-backbone alignment, calculated over all 35 ligand atoms relative to the production-run input structure. All 2,501 frames are shown without smoothing. Selected values at 51.60, 91.80, and 100 ns are annotated. (H) Protein-aligned overlays of ligand poses at 0 ns (green), 90.96 ns (blue), 91.80 ns (orange), 92.12 ns (purple), and 100 ns (red). The initial protein structure is shown in gray. Whole-protein and enlarged local views are presented in the same protein reference frame without ligand self-alignment.

Total radius of gyration (Rg) averaged 2.747 nm over the 100-ns trajectory and ranged from 2.704 to 2.808 nm (Fig 9D). Total solvent-accessible surface area (SASA) averaged 283.3 nm², with a range of 267.4–301.5 nm², and fluctuated without a sustained decrease (Fig 9E). The RMSD–Rg free-energy landscape contained a low-energy region at approximately RMSD 0.25–0.31 nm and Rg 2.74–2.76 nm (Fig 9F). This landscape summarizes conformational sampling along the trajectory.

After correction for periodic-boundary artifacts and protein-backbone alignment, ligand RMSD at 91.80 ns decreased from approximately 16.33 to 1.33 nm. The corrected value was 2.57 nm at 51.60 ns and 2.12 nm at 100 ns (Fig 9G). Protein-aligned overlays showed displacement of the ligand from its initial pose at 100 ns (Fig 9H), consistent with the endpoint ligand RMSD of 2.12 nm.

## Discussion

Kinic-related signature scores differed between healing groups in a cell-type-specific manner, with healing-enriched fibroblasts showing both candidate-gene expression differences and heterogeneity within the population. Inflammatory_Fib and Matrix_Fib cells accounted for larger pooled proportions in healing ulcers. Building on previous descriptions of healing-associated fibroblasts [1,2], these findings localize the exploratory candidate signals within a heterogeneous stromal population and suggest that expression differences and subtype composition should be considered together when interpreting healing-associated tissue profiles.

The opposing score patterns in fibroblasts and myeloid cells argue against assigning a single direction to the association between this transcriptional signature and healing. Such cellular heterogeneity is consistent with the distinct stromal and immune programs described in previous DFU atlases [1,2]. Because the median-score cutoff was applied across all cells, high-versus-low comparisons also incorporate differences in cellular composition. UCell measures relative expression of a gene set within individual cells [10]. Determining how these scores relate to the histone modification characterized by Jiang et al. [3] will require direct biochemical measurements in the relevant cell populations.

LDHB provides a metabolic hypothesis for the fibroblast-associated signal. Work in mammalian cells has linked LDH-associated lactate utilization to mitochondrial metabolism [35]. Here, higher LDHB expression in Healer HE-Fibroblasts and its association with inflammatory, E2F-target, and epithelial–mesenchymal transition gene sets suggest that metabolic activity may accompany cellular activation and tissue remodeling. This interpretation remains provisional because the enrichment analysis was performed in bulk tissue, where changes in cell composition can generate coordinated expression patterns. Testing lactate utilization and fibroblast repair functions after LDHB perturbation would help distinguish a functional contribution from a correlated transcriptional state.

ISG15 offers a link to regulation of the inflammatory environment. Extracellular ISG15 stimulates cytokine secretion through the LFA-1 receptor [36], while intracellular ISG15 stabilizes USP18 and limits type I interferon responses in human cells [37]. The interferon-associated enrichment observed here is compatible with either an activated immune response or compensatory feedback that restrains prolonged signaling. In particular, higher ISG15 expression alone does not identify which of these functions predominates in fibroblasts. Geneformer deletion of ISG15 modestly reduced similarity to the non-Healer reference state, providing a testable hypothesis about its association with fibroblast state. The model does not resolve intracellular versus extracellular mechanisms or establish an effect on wound closure. HAUS1 encodes a component of the augmin complex, which supports centrosome and mitotic spindle integrity [38]. Its higher expression in Healer HE-Fibroblasts could therefore accompany proliferative activity during repair, although proliferation was not measured directly. HAUS1-associated bulk enrichment included inflammatory, IL6–JAK–STAT3, and coagulation programs, connecting its expression with a broader wound-response context. These associations do not establish that HAUS1 controls those pathways. Cell-cycle measurements and gene perturbation within defined fibroblast subtypes would be needed to test that interpretation.

The greater pooled representation of Inflammatory_Fib cells is consistent with the inflammatory features of healing-associated fibroblasts described in the source studies [1,2]. Our subclustering distinguishes variation within that previously annotated population, suggesting that a change in its abundance can also involve a shift in subtype composition. At the tissue level, neutrophil abundance scores correlated with the three candidates, but the between-group abundance comparisons did not survive multiple-testing correction. The two analyses therefore motivate investigation of a shared inflammatory environment while leaving the direction and functional consequences of stromal–immune interactions unresolved.

The compound analyses extend the candidate hypotheses to structural models. The displacement of the LDHB ligand from its initial pose during the single simulated trajectory limits the support that the docking score provides for a persistent interaction. Likewise, docking to the ISG15 C78S model and the HAUS1 fragment requires caution when extrapolating to native full-length proteins. Replicate simulations and direct binding assays would be necessary before proposing these compounds for functional studies of wound repair.

Several limitations define the scope of these findings. First, the bulk cohort was small and the screening thresholds were relaxed during exploratory analysis. None of the three candidates passed adjusted P < 0.05 in that cohort. The single-cell comparisons pooled cells across donors, and metacells could combine cells from different samples. Replication with donor-aware analysis is therefore required. Second, the Kinic-related signature represents an expression-based proxy whose relationship to the modification has not been established in diabetic wounds. Third, the trajectory and virtual perturbations provide computational hypotheses. CytoTRACE informed root selection, and post-perturbation enrichment remains preliminary because the original identifier-conversion records were not retained and GeneRatio denominators remain incompletely reconciled. Fourth, the structural analyses used selected receptor models and one molecular dynamics trajectory, and the study includes no functional validation. Independent cohorts, direct Kinic measurements, and targeted experiments in wound-relevant fibroblasts would address these limitations.

## Conclusions

Kinic-related transcriptional patterns differ between DFU healing groups in a cell-type-specific manner, with healing-enriched fibroblasts showing exploratory candidate-gene expression differences and shifts in pooled subtype composition. These findings connect tissue-level associations to defined wound-cell states and provide hypotheses for understanding heterogeneous healing outcomes.

## Data and code availability

The source RNA-sequencing datasets are publicly available in the Gene Expression Omnibus under accessions GSE231643 [39], GSE165816 [40], and GSE134431 [41]. S1 Data contains the included sample identifiers, selected input and result tables, docking complexes, and corrected molecular dynamics summaries. S2–S4 Data contain the GraphBAN compound inputs and target-specific predictions. The author-generated analysis scripts, processed single-cell objects, simulation input files, and production molecular dynamics trajectory are not included in the supporting files.

## Author contributions

Conceptualization, Zijun Ye, Chenhao Ye and Lin Yang; formal analysis, Chenhao Ye; data curation, Yinghao Wu; visualization, Bohuai Ning; writing—original draft preparation, Zijun Ye; writing—review and editing, Yinghao Wu, Bohuai Ning, Lin Yang and Yiheng Chen; supervision, Yiheng Chen. Zijun Ye and Bohuai Ning contributed to interpretation of the results, Yinghao Wu and Bohuai Ning checked the results and presentation, and Lin Yang contributed to discussion of the results.

## Acknowledgments

We acknowledge the investigators who generated and publicly shared the datasets analyzed in this study.

## List of abbreviations

DEG: differentially expressed gene
DFU: diabetic foot ulcer
ECM: extracellular matrix
GEO: Gene Expression Omnibus
CLS: classification token
GGI: gene–gene interaction
GO: Gene Ontology
GSEA: gene set enrichment analysis
hdWGCNA: high-dimensional weighted gene co-expression network analysis
HE-Fibroblasts: healing-enriched fibroblasts
KEGG: Kyoto Encyclopedia of Genes and Genomes
Kinic: lysine isonicotinylation
ME: module eigengene
NES: normalized enrichment score
Rg: radius of gyration
RMSD: root-mean-square deviation
RMSF: root-mean-square fluctuation
SASA: solvent-accessible surface area
UMAP: Uniform Manifold Approximation and Projection
UMI: unique molecular identifier.

## Supporting information

**S1 Data.** Selected input data and analysis outputs for the diabetic foot ulcer healing study. The archive includes the sample list, bulk count and group tables, the 84-gene input list, selected transcriptomic results, six docking complexes, and molecular dynamics summaries. (ZIP)

**S2 Data.** GraphBAN input compounds and target-specific predictions, part 1 of 3. (ZIP)

**S3 Data.** GraphBAN input compounds and target-specific predictions, part 2 of 3. (ZIP)

**S4 Data.** GraphBAN input compounds and target-specific predictions, part 3 of 3. (ZIP)

